# IF1 connects obesity and insulin resistance through mitochondrial reprogramming in association with ANT2

**DOI:** 10.1101/2020.09.24.311076

**Authors:** Ying Wang, Yaya Guan, Jiaojiao Zhang, Xinyu Cao, Shuang Shen, Genshen Zhong, Xiwen Xong, Yanhong Xu, Xiaoying Zhang, Hui Wang, Jianping Ye

## Abstract

IF1 (ATPIF1) is a nuclear DNA-encoded protein with an activity in the inhibition of catalytic activity of F_1_F_o_-ATP synthase (ATPase), an enzyme for ATP synthesis in mitochondria. A role of IF1 remains unknown in the metabolic disorder in obesity. In this study, IF1 was examined in the diet-induced obese (DIO) mice and a decrease in IF1 protein was observed in several tissues including the skeletal muscle, liver and intestine in the absence of mRNA alteration. Significance of the reduction was investigated in the IF1-KO mice, in which insulin sensitivity was improved in the absence of body weight alteration on Chow diet. On a high fat diet (HFD), the IF1-KO mice gain more body weight as a result of enhanced fat tissue growth. The energy expenditure and locomotion activity were decreased in the KO mice without an alteration in food intake. The increase in insulin sensitivity remained in the obese KO mice. The colon tissue exhibited a resistance to the HFD-induced atrophy with less cell apoptosis and more secretion of GLP-1. Mitochondria exhibited an enhanced ATP production and maximal oxygen consumption without an alteration in the respiratory chain proteins. However, the ATP level was reduced in the fasting condition in the muscle as well as the liver. Mitophagy was enhanced with elevated accumulation of PINK1 and Parkin proteins in the mitochondria. The protein abundance of ADP/ATP translocase 2 (ANT2) was decreased in the inner membrane of mitochondria to account for the reduced apoptosis and enhanced mitophagy. The data suggest that the IF1 reduction in obesity leads to reprogramming of mitochondrial metabolism in a compensatory response to maintain the insulin sensitivity through down-regulation of ANT2 protein.

## INTRODUCTION

Obesity increases the risk of type 2 diabetes through induction of insulin resistance. Energy surplus leads to insulin resistance through suppression of the insulin signaling activity in cells under obesity. The signal of energy surplus remains unknown for the suppression although several candidate molecules have been widely reported, such as ATP, reactive oxygen species (ROS), diglyceride, ceramide, glucose, and free fatty acids, etc. [1; 2]. However, translation of those findings into therapy of type 2 diabetes remains a challenge in the drug discovery field. Inhibition of ATP production represents a mechanism for the therapeutic activity of metformin in the type 2 diabetes patients, which is associated with activation of AMPK pathway in cells [3]. Inhibition of ATP production with chemical uncouplers [4; 5] or berberine [6; 7] is another example for the role of ATP in the mechanism of insulin resistance. These lines of evidence suggest that a surplus in ATP supply is a potential risk factor in the pathogenesis of insulin resistance. However, the possibility remains to be tested in vivo, especially in a transgenic model.

The intracellular ATP level is determined by the balance of production and consumption in cells. The ATP level is maintained at a “set point” level, which varies in cell types for around 5-7 mM in the cardiomyocytes [8]. An increase in ATP consumption leads to more production by the mitochondria to maintain the cytoplasmic ATP hemostasis [9]. Mitochondria activity in ATP production is dependent on the oxidative phosphorylation reaction, which is conducted by the five complexes in the respiratory chain. The F_1_Fo-ATP synthase (ATPase) in the complex V has a well-known activity in the production of ATP through phosphorylation of ADP under conditions of sufficient supply of substrates including oxygen, glucose, fatty acids and amino acids. However, the ATPase also hydrolyzes ATP in the conditions of substrate deficiency in an effort to maintain the mitochondrial membrane potential [10]. The hydrolytic activity of ATPase may prevent ATP surplus in cells in favor of insulin sensitivity. The ATPase activities are controlled by the inhibitor protein IF1, which inhibits both activities of ATPase [10]. In the absence of IF1, the activities of ATPase are enhanced in the production as well as hydrolysis of ATP. An impact of such reprogramming of ATPase activities in insulin sensitivity remains to be tested after IF1 inhibition, especially following the enhanced hydrolytic activity of ATPase. To address this issue, we conducted this study in IF1-KO mice, which included both male and female mice.

We examined the IF1 in the DIO mice and observed reduction in the protein in the muscle, liver and intestine. A role of the reduction was examined in the IF1-KO mice, in which obesity was enhanced without an extra impairment in insulin sensitivity. The phenotype was associated with mitochondrial reprogramming in energy metabolism and quality control, which was observed with a reduction in ANT2 protein.

## RESEARCH DESIGN AND METHODS

### Generation of IF1-KO mice and DIO mice

All of the animal procedures were approved by the Institutional Animal Care and Use Committee (IACUC) of the Shanghai Jiao Tong University. The IF1 (ATPIF1) knockout mouse was generated in C57BL/6 mice with the CRISPR/Cas9 gene editing method with sgRNA PAM sequences targeting ATPIF1: SgRNA-1 GCAGTCGGATAGCATGGATACGG and SgRNA-2 GGCTCCACCAGCTTCTCGGATGG. For identification of IF1^-/-^ mice, PCR was conducted to amplify the IF1 gene using the forward primer 5-CATCAGCCTTGGAATTCTGC-3 and the reverse primer 5-CTTCGTCTCGGACTCGGTAG-3. The agarose electrophoresis was performed to determine the genotype, and the amplified PCR product was used in gene sequencing with ABI-3730XL to confirm the genotype. The IF1 protein was examined to confirm the gene knockout. The mouse housing environment includes a 12:12-hr light-dark cycle, constant room temperature (22–24°C), free access to water and diet. The mice were fed on Chow diet (Fat content ≥ 4% in weight, P1101F-25, Shanghai Pluteng Biotechnology Co., Ltd., China) or the high fat diet (HFD, Fat = 40% in kcal, D12108C, Jiangsu Synergy, China). HFD feeding was started at 8 wks in age to generate the diet-induced obese (DIO) model. Age- and gender-matched wild type mice were used in the control for the IF1-KO mice.

### Body weight, body composition and food intake

Body weight of mice was measured weekly. The body composition was determined using the quantitative nuclear magnetic resonance (NMR) (Minispec Mn10 NMR scanner, Brucker, Milton, ON, Canada) in the conscious and unrestrained mice, which was placed individually in a small tube for the NMR analyzer with 1 min in the assay. Food intake was measured manually or with the metabolic chamber individually on HFD. A mean value of daily food intake was determined over 3 days. Unit of food intake was g/mouse/day.

### Energy expenditure and physical activity

Energy metabolism was monitored in the mice after 4 wks on HFD, using the 20-cage Promethion-C continuous, parallel metabolic phenotyping system (Sable Systems International, Las Vegas, USA). Mice were kept in the metabolic chamber individually for 6 days. The oxygen consumption (VO_2_), carbon dioxide production (VCO_2_), spontaneous physical activity and food intake were recorded daily for 5 days. The data on day 5 were used in the calculation of energy expenditure (EE: kcal/kg/h) and physical activity. The calculation was performed with the formula EE = [3.815 + 1.232 × VCO_2_/VO_2_] × VO_2_ × 0.001 [11]. Energy expenditure data was normalized with the body lean mass.

### ITT and GTT

Intraperitoneal insulin tolerance test (ITT) was performed in mice (9 wk on Chow diet and 14 wk on HFD) by peritoneal insulin injection (0.75 U/kg body weight, I9278, Sigma) after 8 hr fasting. In DIO mice, oral glucose tolerance test (oGTT) and intraperitoneal glucose tolerance test (ipGTT) were performed in the mice (12-16 wks on HFD) by administration of glucose (2 g/kg body weight) after overnight fasting. Blood glucose was measured in the tail vein blood at 0, 15, 30, 60, 90 and 120□ min using the Accu-CHEK Advantage blood glucose meter (Accu-CHEK; Indianapolis, IN). Data were expressed in blood glucose concentration (mmol/l, mM).

### Western blotting

Tissues (liver, muscle and colon) were collected from the mice after 16 wks on HFD and examined for IF1 protein, insulin signaling, apoptosis and mitophagy. The whole cell lysates were prepared from the tissue in Western blotting according to the protocols described elsewhere [11]. Antibodies to ATPIF1 (ab110277), VDAC1 (ab14734), PINK1 (ab186303), Parkin (ab77924), total OXPHOS Rodent (ab110413), P62 (ab56416), β-actin (ab8224) and GAPDH (ab7291) were obtained from Abcam (Cambridge, MA, USA). Antibodies to Agt7 (8558s), ANT2 (14671S), Caspase 3 (9662S), cleaved caspase 3 (9664S), Akt (C67E7) and p-Akt (T308, C31E5E) were purchased from the Cell Signaling Technology (Boston, USA). GAPDH and β-actin were used as the internal controls. Western blot images were quantified with the ImageJ software, whose value is expressed by integrated optical density (IOD).

### Quantitative real time PCR (qRT-PCR)

The total mRNA was extracted from the tissue using TRIzol reagent according to the manufacturer’s protocol (TAKARA, Japan). The qRT-PCR assay was performed with a kit of TB Green® Premix Ex Taq™ II (Tli RNaseH Plus) (TAKARA, Japan) using the ABI-3730XL machine. The target mRNA was normalized to the ribosome 18S RNA of the endogenous control. Primers and probes were purchased from TaKaRa (RR820A, Nojihigashi, Kusatsu, Shiga, Japan). Sequence of GLP-1 (Gcg) green primers include forward primer 5-TTACTTTGTGGCTGGATTGCTT-3, and reverse primer 5-AGTGGCGTTTGTCTTCATTCA-3. The primers of IF1 are forward: 5-ACGGGCGCTGGCTCCATCC-3; reverse: 5-TGGCGTTCAATTTGCTTCTG-3.

### Insulin and GLP-1 assay

Insulin and GLP-1 were tested in the plasma of ocular vein blood in mice at 16 wks on HFD. Plasma insulin was measured using an insulin ELISA Kit (90080, Crystal Chem, Downers Grove, USA). The blood for GLP-1 assay was collected with a heparin tubes containing dipeptidyl peptidase IV inhibitor (10 μl/mL), which was conducted at 15 min after the glucose administration. The active form of GLP-1 was measured with the GLP-1 ELISA Kit (E-EL-M0090c, Elabscience, Wuhan, China).

### HE staining and transmission electron microscopy

The fresh intestinal tissues were fixed in 4% paraformaldehyde for 24 h in the HE staining, and the electron microscope fixative at 4 □ for 2 h in the electron microscopy study. The samples were prepared by the Wuhan Servicebio technology Co., LTD.

### Mitochondrial and ATP assays

The mitochondria were isolated from the fresh muscle tissue of mouse using a protocol modified from our early study and used in the functional assay with the Agilent Seahorse XF24 Analyzer equipment (Agilent,Santa Clara, California, USA) [12]. When the muscle tissue was used in the mitochondrial assay, the fresh tissue was washed with the Agilent Seahorse XF-DMEM medium (the assay medium) containing 2 mM L-glutamine, 1 mM sodium pyruvate and 10 mM glucose to remove the non-fatty substances. A piece of tissue (2 × 2 × 1 mm) was placed in each well of the Agilent Seahorse XF24 islet capture microplate (103518-100,Agilent,Santa Clara, California, USA) and covered with an islet capture screen to allow free perfusion while minimizing tissue movement. The oxygen consumption rate (OCR) was determined in the Agilent Seahorse XF Complete Assay Medium (500 μL) with the Agilent Seahorse XF24 Analyzer. Following agents were used in the stimulation or inhibition of mitochondrial activity in the assay: ADP (4 mM, A5285, Sigma), Oligomycin (Oligo, 1 μM, 495455, Sigma), FCCP (2 μM, C2920, Sigma), and Antimycin A (AA, 0.5 μM, 2247-10, Biovision). The tissue ATP concentration was determined in the homogenization of fresh tissues using the EnzyLightTM ATP Assay Kit (EATP-100, BioAssay Systems, Northern California, USA).

### Statistical Analysis

Statistical analysis was performed using two-tailed, unpaired Student’s t-test in the study. *p* < 0.05 was considered significant. Results are presented as mean ± SD.

## RESULTS

### IF1 protein is decreased in multiple tissues of DIO mice

IF1 is a mitochondrial protein that associates with F_1_F_o_-ATPase to inhibit the catalytic activities. There was no systematic study of distribution of IF1 protein in the mouse tissues when this project was initiated. We addressed this issue by examining IF1 protein in 9 major tissues of mice with normalization to the internal control of GAPDH protein. The protein levels from high to low were in the following order: brown fat, heart, kidney, liver, spleen, lung, colon, brain, and skeletal muscle (Fig. 1A). An impact of obesity in the IF1 protein was investigated in DIO mice to understand the relationship of IF1 and obesity. The IF1 protein was examined in the insulin sensitive tissues including brown and white fat, skeletal muscle and liver tissues. A dramatic decrease was observed in the IF1 protein in all tissues (Fig. 1, B-D). In addition, the IF1 protein was reduced by obesity in the colon (Fig. 1E). mRNA of IF1 was not altered in the DIO mice (Fig. 1E). The results suggest that IF1 protein was expressed in all tissues and the protein abundance was reduced by obesity.

**Fig. 1.**
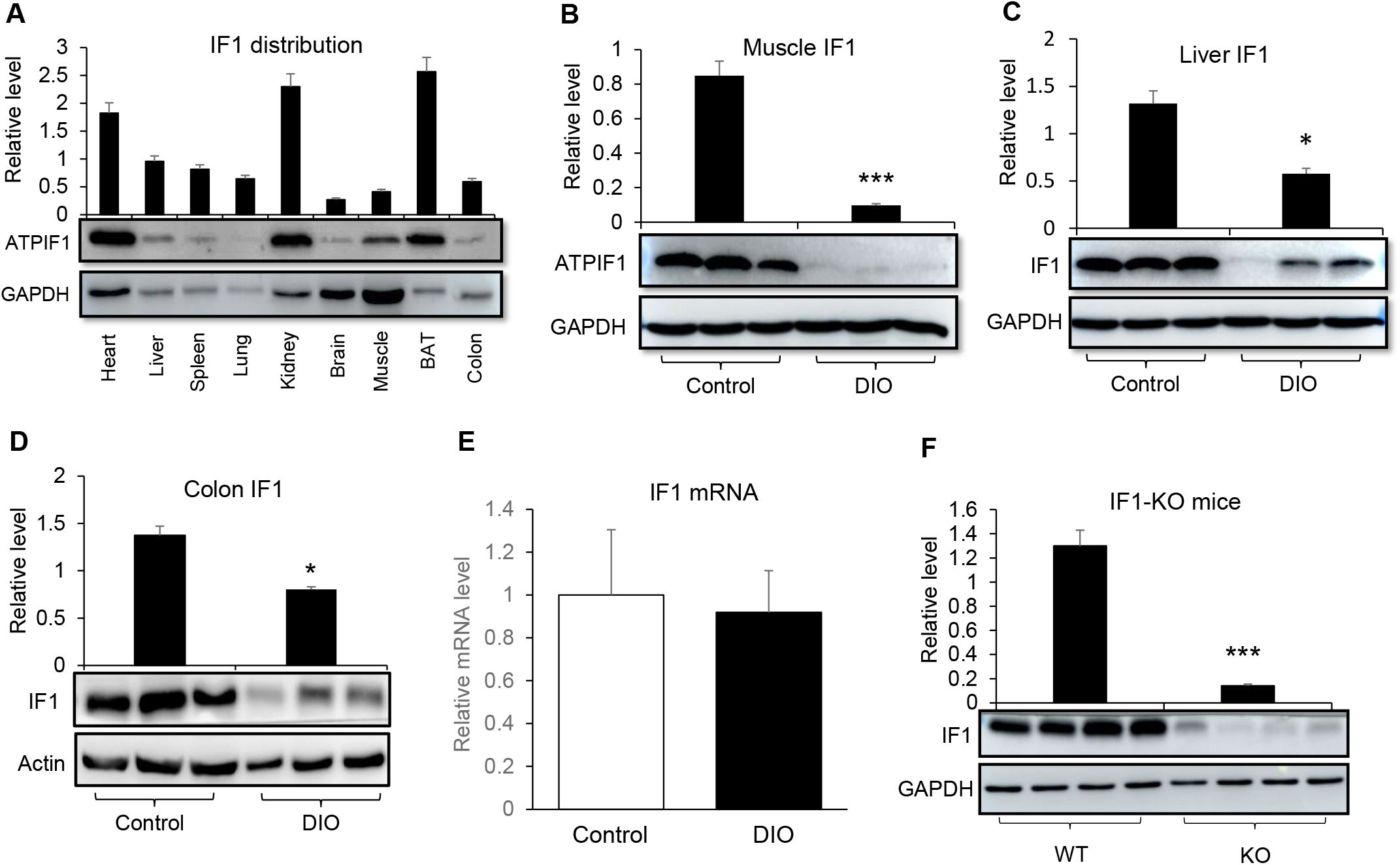
Reduction of IF1 protein in DIO mice. A. Distribution of IF1 protein in the mouse tissues including heart, liver, spleen, lung, kidney, brain, muscle, BAT and colon; B. Abundance of IF1 protein in the skeletal muscle of lean and obese mice; C. Abundance of IF1 protein in the liver tissue of lean and obese mice; D. Abundance of IF1 protein in the colon tissue of lean and obese mice; E. mRNA of IF1 in the muscle and liver tissues (n=6). F. Validation of IF1 knockout by protein assay. The assay was conducted in the skeletal muscle (n=4). The data in bar figure represents mean ± SD. * p<0.05; *** p<0.001.

### ATP production and mitochondrial spare capacity are enhanced in mitochondria of IF1-KO mice

To understand the biological significance of IF1 reduction, we generated the IF1-KO mice with successful deletion of IF1 protein in the mice (Fig. 2A). The mitochondrial oxygen consumption rate (OCR) was examined in the skeletal muscle of KO mice to evaluate the impact of IF1 inactivation in mitochondrial function. The study was conducted in isolated mitochondria in response to challenges of ADP and uncoupler FCCP using the Seahorse equipment. The mitochondria exhibited an increase in the respiration as indicated by the elevated OCR at the basal and challenged conditions (Fig. 2, B and C). The protein abundance was not significantly altered in the respiratory chain of mitochondria in the IF1-KO mice (Fig. 2D). The data suggest that in the absence of IF1, the production of ATP upon ADP challenge is enhanced for the elevated activity of ATPase. The alteration leads to an increase in the mitochondrial spare capacity as indicated by the elevation in the maximal OCR upon FCCP challenge.

**Fig. 2.**
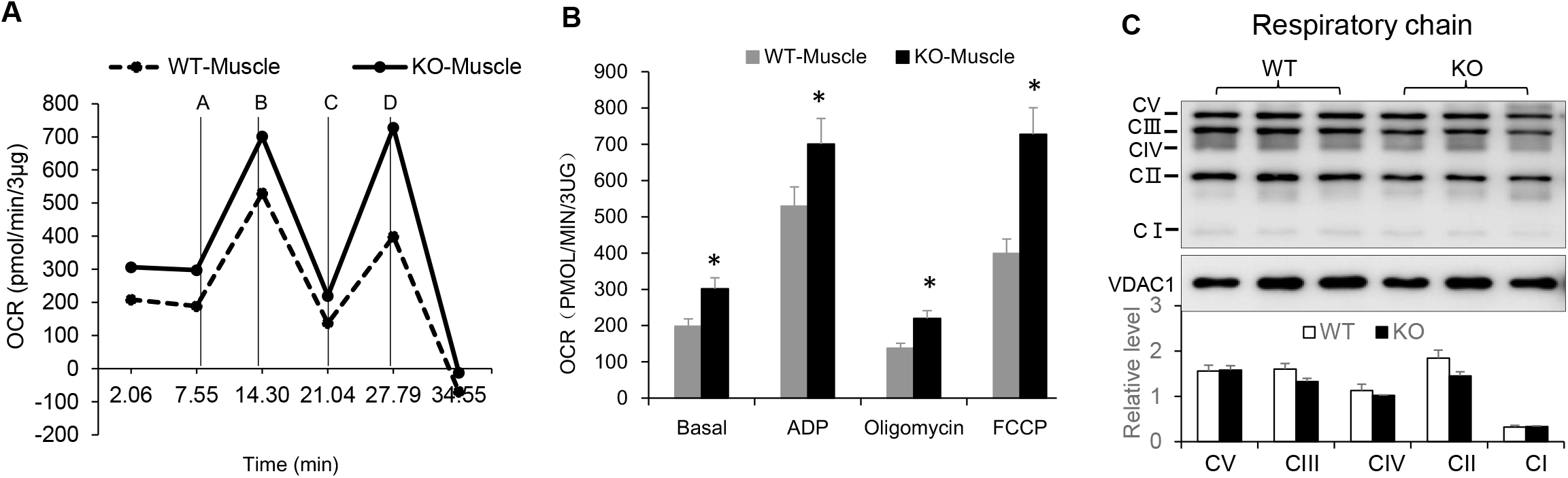
Mitochondrial function was enhanced in the IF1-KO mice. A. Mitochondrial function determined by oxygen consumption rate (OCR). The test was conducted in mitochondria isolated from the skeletal muscle tissue of male mice at 8 weeks in age. The injected agents were A: ADP; B: Oligomycin; C: FCCP; D: Antimycin A. B. ATP production by mitochondria. The ATP production was calculated with the OCR induced by ADP in the reaction. C. Proteins in the respiratory chain of mitochondria. The representative proteins were examined in the muscle tissue by Western blotting. The data represents mean ± SD (n=3). * p<0.05.

### Insulin sensitivity is enhanced in IF1-KO mice on Chow diet

The IF1 protein reduction was associated with insulin resistance in the DIO mice. However, the cause/effect relationship was unknown for the two events. To address this issue, insulin sensitivity was examined in the IF1-KO mice on Chow diet in both genders. The insulin tolerance test was conducted with intraperitoneal injection of insulin (ipITT). The IF1-KO mice exhibited a significant increase in insulin sensitivity as indicated by more reduction in the blood glucose in the KO mice over the WT mice (Fig. 3, A and B). There was no significant difference in the body weight between the KO and WT mice (Fig. 3, C and D). The data suggest that inhibition of IF1 activity by gene knockout led to an increase in insulin sensitivity in the IF1-KO mice.

**Fig. 3.**
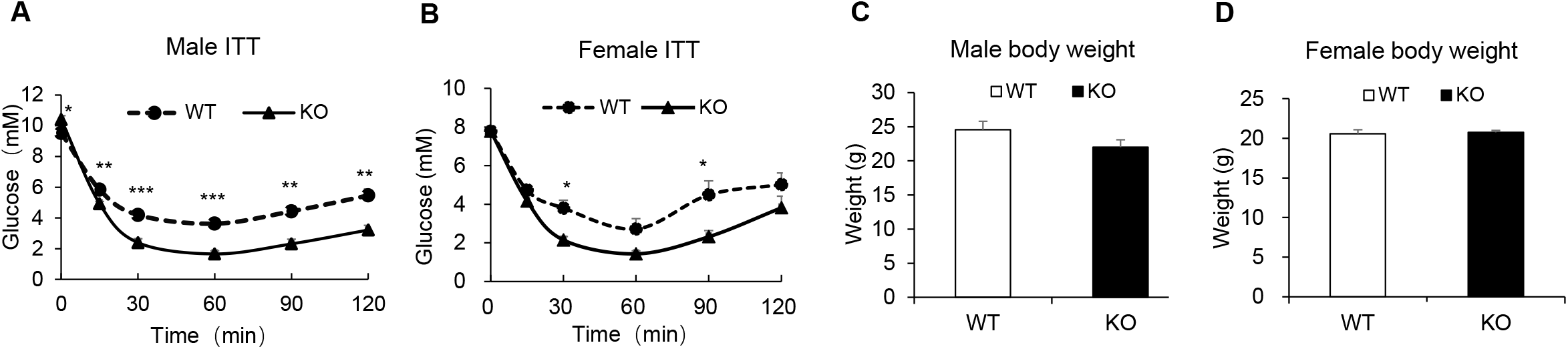
Insulin sensitivity of the IF1-KO mice on Chow diet. Insulin tolerance test and body weight were determined in male and female mice on Chow diet at 8 weeks of age. A. ITT in male mice; B. ITT in female mice. C. Male mice body weight. D. Female mice body weight. The data represents mean ± SD (n=6). * p<0.05; ** p<0.01.

### IF1-KO mice gains more fat tissue on HFD

Obesity was induced in the IF1-KO mice with HFD feeding to test the mice responses to the energy surplus. The IF1-KO mice gained more body weight than the WT mice in both gender (Fig. 4, A and B). The initial body weight was identical in the female mice, but modestly higher in the male mice before the feeding. The KO mice exhibited a higher gain in the body weight than WT mice throughout the study. A difference in the fat accumulation contributed to the extra weight gain in the KO mice as the total body fat mass, epididymal fat pad, and liver size were all higher in the KO mice (Fig. 4, C and D). Energy expenditure were examined using the rodent metabolic chamber. A reduction was observed in the IF1-KO mice at the day- and night-time (Fig. 4E). A reduction in the physical activity was observed, but a significance was only obtained at the nighttime (Fig. 4F). Food intake was not significantly altered in the mice (data not shown). These data suggest that inactivation of IF1 gene led to a reduction in energy expenditure for an increased risk of obesity in the IF1-KO mice.

**Fig. 4.**
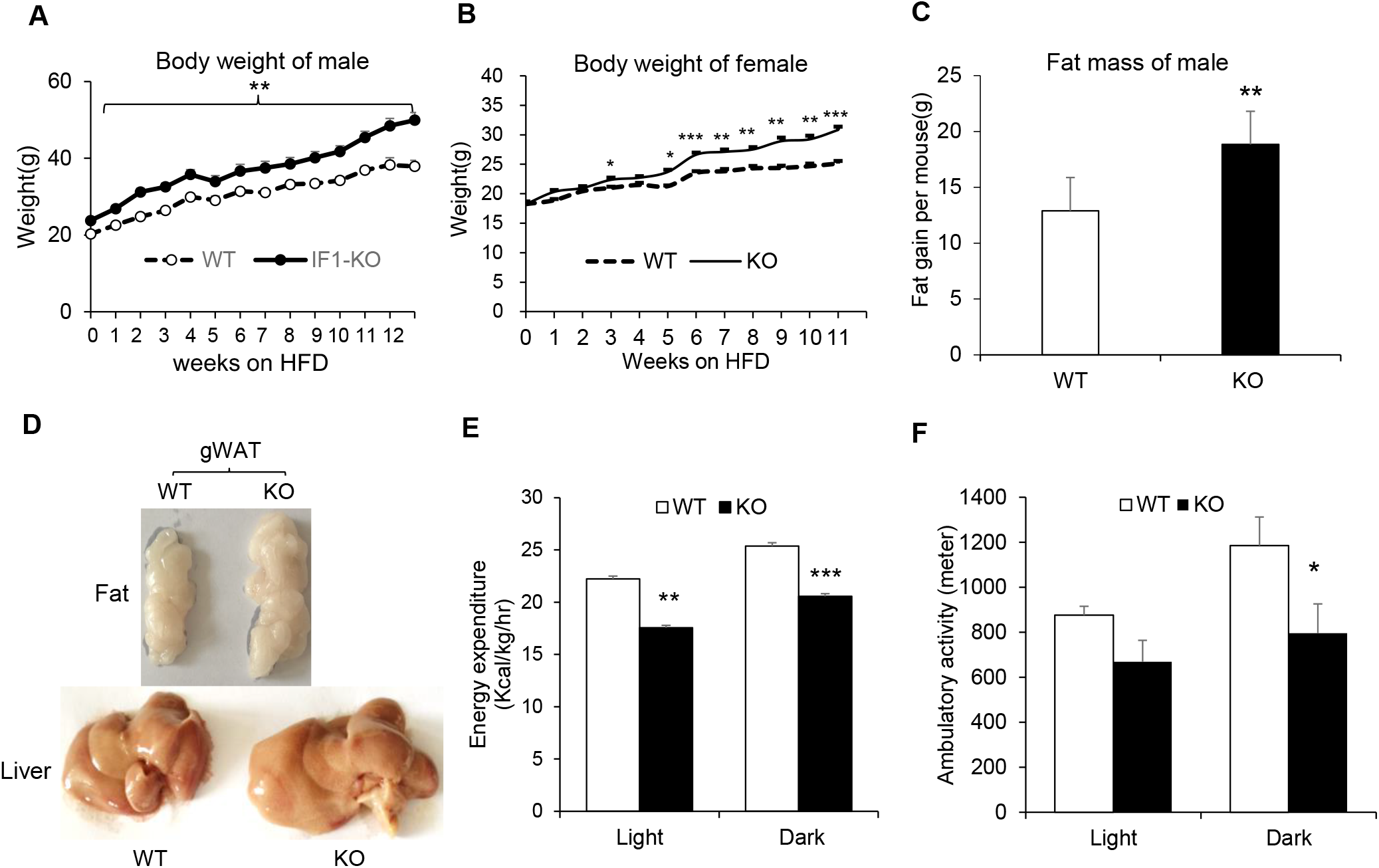
Obesity was enhanced in the IF1-KO mice. A. Male mice body weight on HFD (n=9). B. Female mice body weight on HFD (n = 8-12); C. Fat mass of male mice on HFD (n=9); D. Representative images of epididymal fat and liver of mice on HFD;E. Energy expenditure based on oxygen consumption rate in the female IF1-KO mice after 4 weeks on HFD (n=8). F. Physical ambulatory activity in the female IF1-KO mice after 4 weeks on HFD (n=8); The data represents mean ± SD. * p<0.05; ** p<0.01.

### Heat production is increased in the IF1-KO mice

The reduction in energy expenditure was observed in the female KO mice in above experiment. The same experiment was conducted in the male mice. A similar reduction was observed in the energy expenditure and physical activity in the male mice (Fig. 5, A-D). The reduction was observed at the nighttime, which corresponded to the reduced oxygen consumption rate at night. However, food intake was not decreased in the presence of lower physical activity (Data not shown). Interestingly, heat production was significantly increased in the male IF1-KO mice (Fig. 5, E and F). These results suggest that the reduction in physical activities contributes to the low energy expenditure in the KO mice. In the energy expenditure, more heat was produced in the KO mice.

**Fig. 5.**
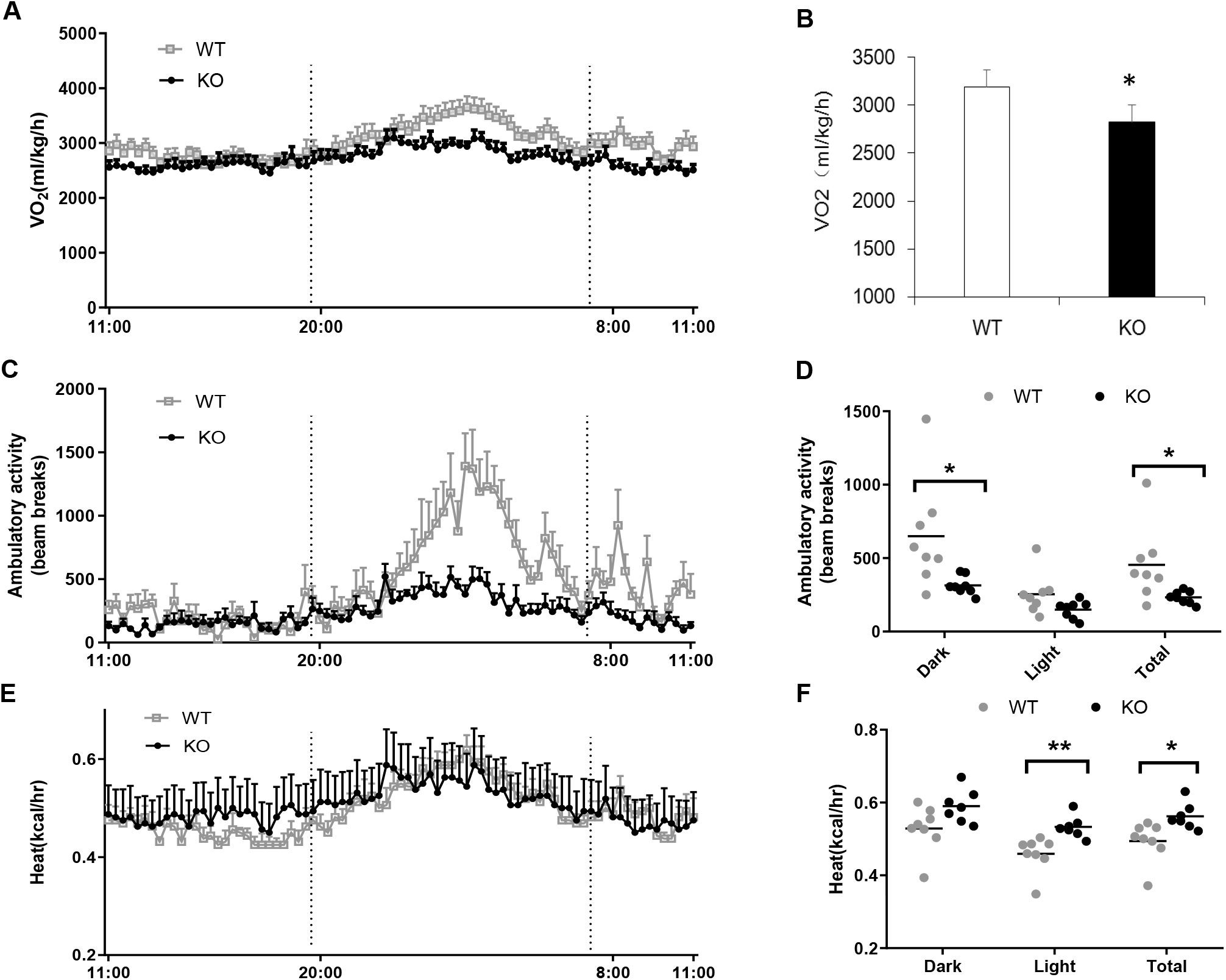
Heat production is increased in the IF1-KO mice. A. OCR of male IF1-KO mice on HFD (n=8). B. OCR in male IF1-KO mice at nighttime on HFD; C. Physical ambulatory activity of male mice on HFD; D. Mean value of ambulatory activity in daytime and nighttime; E. Heat production of male IF1-KO mice on HFD; F. Mean value of heat production of male IF1-KO mice in daytime and nighttime. The data represents mean ± SD (n=8). * p<0.05; ** p<0.01.

### Insulin resistance is disassociated with obesity in the IF1-KO mice

In the IF1-KO mice, insulin sensitivity was examined to determine the impact of obesity in the glucose metabolism. The study was conducted in several tests including the intraperitoneal glucose tolerance (ipGTT), insulin tolerance (ipITT) and insulin signaling tests. The KO mice exhibited an improved insulin sensitivity in all the assays (Fig. 6, A-D). In the GTT assay, the blood glucose was increased much less in the KO mice (Fig. 6A). In the ITT assay, the blood glucose was decreased more in the KO mice (Fig. 6B). The insulin signaling pathway was examined in the skeletal muscle and liver tissues of KO mice with Akt phosphorylation. An increase was observed in both tissues (Fig. 6, C and D). The ATP level was decreased in the muscle and liver of IF1-KO mice in the fasting condition (Fig. 6, E and F). This group of data suggests that insulin sensitivity was not preserved in the KO mice in the presence of increased adiposity. The phenotype suggests a disassociation of insulin resistance and obesity in the KO mice.

**Fig. 6.**
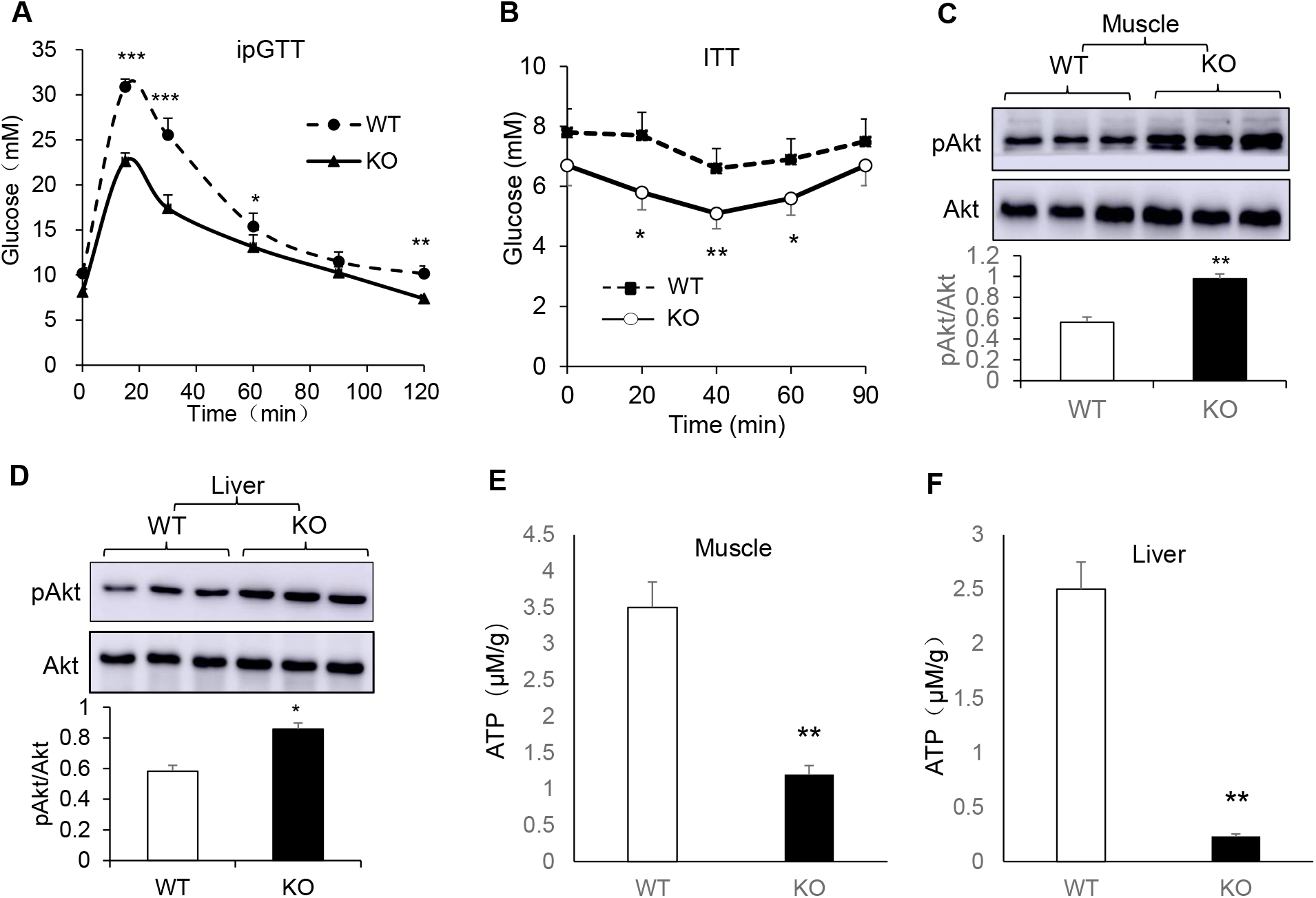
Insulin sensitivity was enhanced in the IF1-KO mice. The female mice were subject to the glucose tolerance test (GTT), insulin tolerance test (ITT), insulin signaling test and ATP test sequentially between 12 – 16 weeks on HFD. The intraperitoneal glucose tolerance test was conducted with glucose (2 g/kg) after overnight fasting at 12 weeks on HFD. The blood glucose was determined using the glucose meter at 5 time points (n=8 -12). B. Insulin tolerance test (ITT). The same mice were injected with insulin (0.75 U/kg) intraperitoneally at 14 weeks on HFD after 8 hr fasting. C. Insulin signaling in muscle. The signal of pAkt was detected in the muscle of mice at 30 min after insulin injection (0.75 U/kg body weight) at 16 weeks on HFD (n=3); D. Insulin signaling in the liver. The test was conducted in the liver under the same condition as the muscle assay (n=3). E. ATP level in muscle. The tissue ATP level was determined in the skeletal muscle in the same samples (n=6). F. ATP level in the liver. The ATP level was determined in the liver tissue in the same samples (n=6). The data represents mean ± SD. * p<0.05; ** p<0.01.

### Plasma GLP-1 is elevated in the IF1-KO mice

Above data suggest that glucose metabolism was reprogrammed in the IF1-KO mice to preserve insulin sensitivity. The impact of mitochondrial reprogramming in GLP-1 activity was investigated to understand the contribution of endocrine activity to the glucose homeostasis. GLP-1 acts in multiple tissues in the regulation of blood glucose in the control of type 2 diabetes. GLP-1 activity was examined in the KO mice on HFD through an oral glucose tolerance test (OGTT), which induces GLP-1 secretion in the intestine. OGTT was significantly improved in the IF1-KO mice (Fig. 7A). Plasma GLP-1 level was examined during the OGTT test, and an increase was observed in the IF1-KO mice (Fig. 7B). Expression of GLP-1 was examined in mRNA and a significant elevation was observed in the large intestine of the IF1-KO mice (Fig. 7C). In the DIO mice, GLP-1 secretion is decreased in the large intestine from a tissue atrophy following mitochondrial degeneration [12]. The colon tissues were examined in the IF1-KO mice to understand the mechanism of improved GLP-1 secretion. The thickness of intestine wall and mucosa layer were measured in the evaluation of intestine atrophy after H&E staining. An increase in the thickness was observed in the IF1-KO mice with abundant smooth muscle and mucosa content (Fig. 7D). These data suggest that GLP-1 secretion was improved in the intestine of IF1-KO mice, which may contribute to the disassociation of insulin resistance and obesity in the obese IF1-KO mice.

**Fig. 7.**
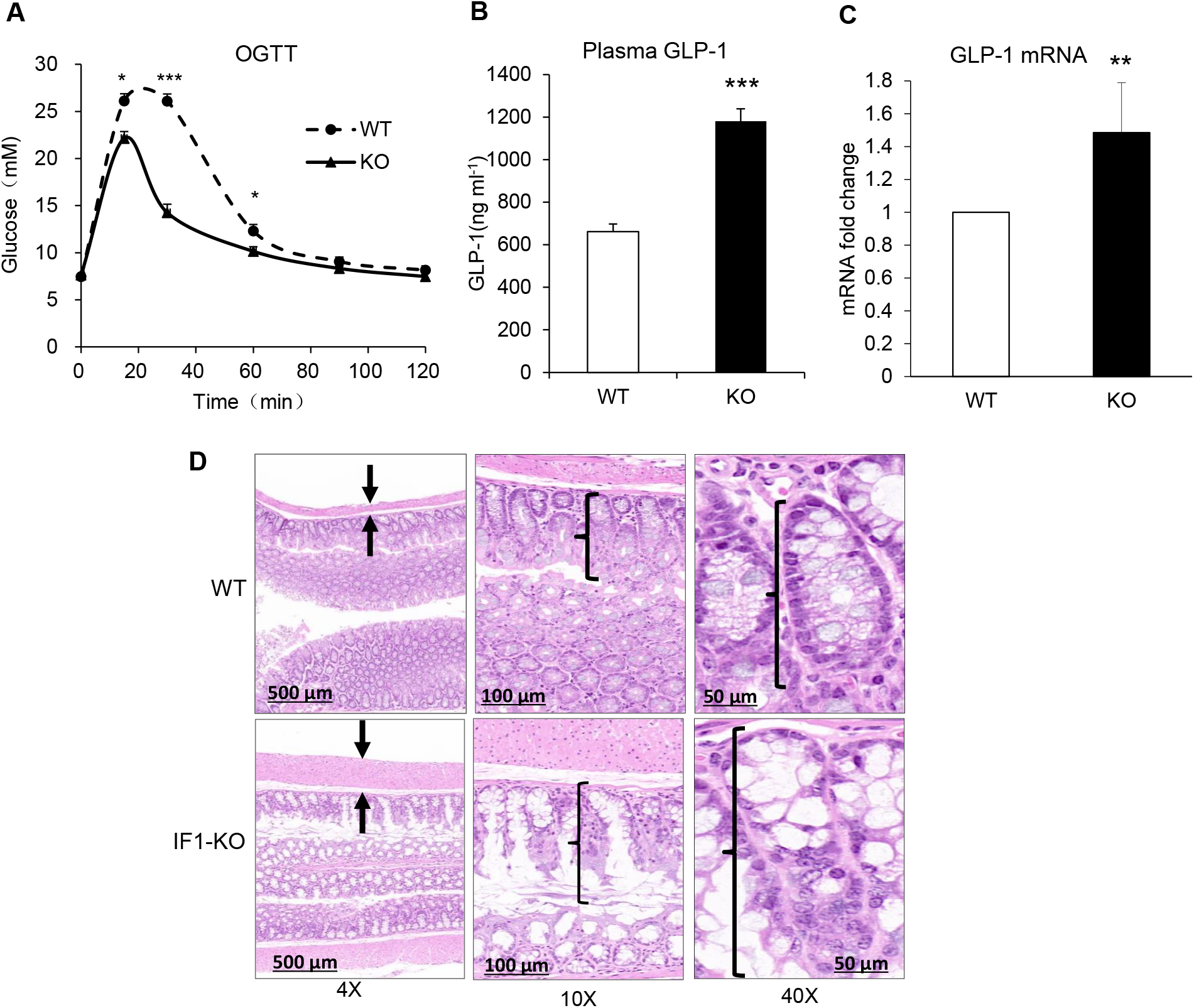
GLP-1 was increased in the IF1-KO mice. A. Oral glucose tolerance test (oGTT). The test was conducted in the female mice after overnight fasting at 15 weeks on HFD by oral administration of glucose (2g/kg) (n=8-12); B. Plasma GLP-1 level. The test was conducted at 15 min post oral glucose administration using an enzyme-linked immunosorbent assay kit (p=0.000339) (n=5); C. GLP-1 mRNA in the colon. mRNA was determined by qRT-PCR in the colon tissue of mice at 16 weeks on HFD (n=5). D. The anti-atrophy activity of colon. The colon tissue slide was subjected to H&E staining and observed under the microscope. The tissue thickness is indicated by the arrows or markers. The data represents mean ± SD. * p<0.05; ** p<0.01; *** p<0.001.

### Mitochondria exhibits an improved quality control in the IF1-KO mice

Villi and mitochondrial morphology were examined in the colonic epithelial cells under the electron microscope. The villi were heathier in the IF1-KO mice with persistent length versus the sickness type of villi of different length in the WT mice (Fig. 8A). In the super structure, mitochondria had smaller size and higher matrix density in the epithelial cells of the IF1-KO mice (Fig. 8A), which are signs of healthy mitochondria. In contrast, mitochondria of the WT mice had a larger size and less crista in the parameters of mitochondrial degeneration. Mitochondria control tissue atrophy through an impact in apoptosis. The mitochondrial alteration may reduce tissue atrophy through downregulation of apoptosis in the IF1-KO mice. To test the possibility, the cleaved caspase 3 (c-Cas3) signal was examined to determine apoptosis, and a reduction was observed in the large intestine of IF1-KO mice (Fig. 8B). Adenine nucleotide translocase 2 (ANT2), a ADP/ATP translocase in the mitochondrial inner membrane, contributes to apoptosis by mediating proton leak in the collapse of mitochondrial potential [13]. A decrease in ANT2 activity leads to less proton leak in the inhibition of cell apoptosis [14; 15]. ANT2 protein was significantly reduced in the colon tissue of IF1-KO mice (Fig. 8B). Mitophagy markers, PINK1 and Parkin, were examined to determine the mechanism of mitochondrial activity in apoptosis. Both signals were increased in the IF1-KO mice colon (Fig. 8C). The autophagy markers including p62 and Atg7 were not changed in the same condition. Under the electronic microscope, mitophagy was observed with mitochondrial structure in the autophagosome (Fig. 8B), in which more mitochondria were found in the autophagosome of IF1-KO mice as pointed by the white color arrows. These data suggest that the reduction in tissue atrophy may be a result of suppression of cell apoptosis following the decrease in the ANT2 protein together with the increased mitophagy activity.

**Fig. 8.**
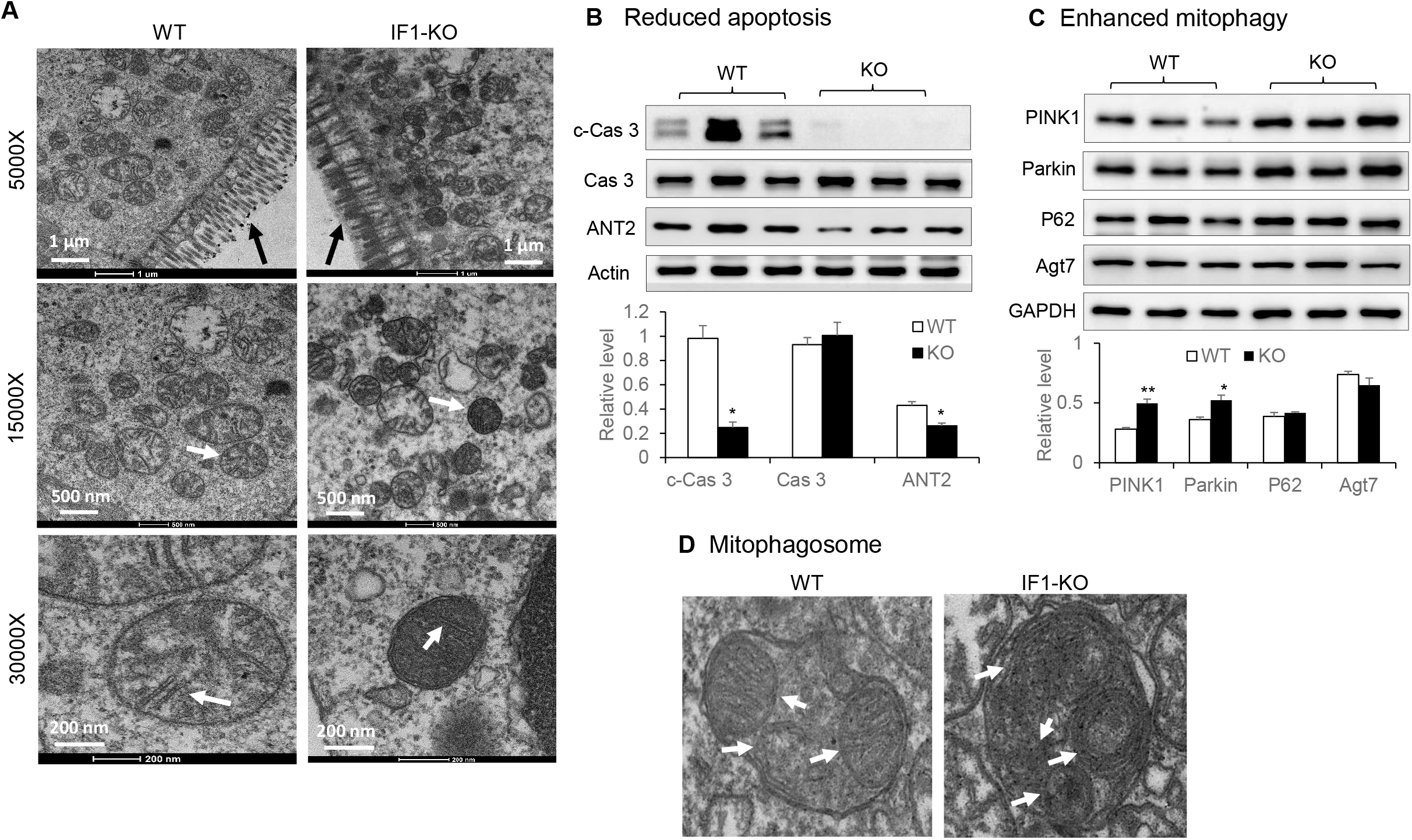
Mitophagy was enhanced in IF1-KO mice. A. Villi and mitochondria were healthier in the colonic epithelial cells of IF1-KO mice. The colon tissue was collected from the female mice on HFD for 16 weeks and examined under the electronic microscope; B. Reduced cell apoptosis in the colonic tissue of IF1-KO mice. The representative proteins of apoptosis markers were examined by Western blotting. C. Mitophagy markers in the colon tissues. The markers were determined by Western blotting. D. Electronic microscope image of mitophagy. The bar figure presents data of mean ± SD (n=3). *p<0.05, **p<0.01.

## DISCUSSION

The study demonstrates that the IF1 protein is expressed in all tissues, and the abundance is reduced by obesity. There was no information about regulation of IF1 protein in the normal tissues in the literature except tumor tissue [16; 17], in which IF1 protein is increased to promote glycolysis in certain types of tumor [16; 18]. In this study, the IF1 protein was identified in all tissues with the highest level in the heart muscle, kidney and brown fat of mice, which is consistent with the mRNA expression level in those tissues in a recent study [19]. There is very limited information about regulation of IF1 protein by posttranslational modification. IF1 is subject to phosphorylation by PKA in the functional inhibition and modification by proton for activation in the acidic condition of mitochondria [19; 20]. However, an impact of those modifications in the protein stability remains unknown [17]. Mitochondrial proteins are studied in the human skeletal muscle of obese patients with the proteomics technology [21]. However, the study does not identify a change in the IF1 protein. The IF1 protein was examined by Western blotting in a study of the skeletal muscle, and an increase was observed in the obese subjects [22], which is opposite to ours. The cause for the difference is unknown. The difference in the model systems may play a role. As mRNA of IF1 was not changed, our data suggests that the reduction in IF-1 protein is a result of translational regulation or post-translational modification, which is supported by the feature of mitochondrial proteins [21; 23]. The exact mechanism remains to be investigated for the IF1 protein reduction.

Our study reveals that a metabolic reprogramming is induced in mitochondria by IF1 inactivation. The biological significance of the IF1 reduction was investigated in mitochondria of skeletal muscle, liver and intestine. The ATP production and the maximal oxygen consumption were all elevated in the isolated mitochondria of the IF1-KO mice. The increased ATP production reflects an elevation in the synthetic activity of ATPase enzyme in the absence of IF1 [10], which associates with the increase in the maximal oxygen consumption in response to the uncoupler FCCP. In the fasting condition, ATP level was decreased in the skeletal muscle and liver of IF1-KO mice, which is consistent with the increased hydrolytic activity of ATPase in the absence of IF1 [10]. The metabolic reprogramming was not associated with a change in the protein abundance of the respiratory chain. These data suggest that the elevation in catalytic activities of ATPase is a key in the metabolic reprogramming in mitochondria.

The study demonstrates that the mitochondrial reprograming increased the risk of obesity in the IF1-KO mice, which was observed in both male and female mice. There are two potential mechanisms for this obesity phenotype. The first is reduction in the locomotion activity in the IF1-KO mice for the less energy expenditure as indicated by the fall in oxygen consumption, which was observed in the mice on HFD in the metabolic chamber. The reduction did not lead to a decrease in the food intake in the IF1-KO mice. The second mechanism is an enhancement of adipocyte differentiation that was observed in vitro for an increased accumulation of triglyceride and enlarged lipid droplets in differentiated adipocytes from preadipocytes of the IF1-KO mice [24]. The obesity is supported by the phenotypes of IF1 overexpression mice, in which ROS production is enhanced [25] and inflammatory response is elevated [26]. ROS and inflammation promote energy expenditure in favor of suppression of obesity [27].

The mitochondrial reprograming disassociated obesity and insulin resistance in the IF1-KO mice of both genders. In the WT mice, obesity is positively associated with insulin resistance. However, this rule was broken in the IF1-KO mice on HFD, which exhibited more adiposity without extra impairment in insulin sensitivity. On the Chow diet, the IF1-KO mice exhibited no difference in the body weight, but a better insulin sensitivity over the WT mice. The IF1-KO mice were normal in the growth, development and reproduction on the Chow diet, which is consistent with that in a study by Nakamura [28]. Obesity and insulin sensitivity were not examined in the IF1-KO mice in Nakamura’s study. Our data suggest that IF1 may inhibit insulin sensitivity in the physiological condition, which was abolished in the IF1-KO mice.

The mitochondrial reprograming increased GLP-1 secretion, which provides an endocrine mechanism for the enhanced glucose metabolism in the IF1-KO mice. There is no report about the relationship of IF1 and GLP-1 in the literature. In the WT mice, GLP-1 secretion is reduced by HFD through a tissue atrophy in the intestine [12], which is associated with mitochondrial degeneration and cell apoptosis [29]. In the IF1-KO mice, both pathological alterations were prevented by the mitochondrial reprogramming. Consistently, IF1 inhibition by gene knockdown led to inhibition of apoptosis through stabilization of mitochondrial potential [30]. However, the mitochondrial membrane permeability was not investigated in that study. We addressed this issue by testing ANT2 in the IF1-KO mice.

The mitochondrial reprograming was associated with ANT2 reduction in the IF1-KO mice. ANT2 is a transporter protein located in the inner membrane of mitochondria to mediate the import of ADP from the cytoplasm and export of ATP from mitochondria [31]. ANT2 belongs to the ANT family whose activity is regulated by the Bcl family proteins in the control of apoptosis [13]. However, the exact action remains unknown for the ANT proteins until that ANT1 was found to transport proton in a study of mitochondrial uncoupling activity [32]. An increase in the ANT activity makes the cells more susceptible to apoptosis for a risk of the mitochondrial potential collapse [14]. It was unknown how IF1 regulates ANT in mitochondria. In current study, the ANT2 activity was reduced in the IF1-KO mice, which provides a mechanism for the resistance to apoptosis in support of the GLP-1 secretion function.

The ANT2 reduction may contribute to the enhanced insulin action through mitophagy in the IF1-KO mice. ANT2 gene knockout improves insulin sensitivity in two studies of fat and liver in mice [33; 34]. The exact mechanism remains to be investigated for the enhanced insulin action although a reduction in inflammation was proposed [33]. Mitophagy removes the senescent and damaged mitochondria through self-digestion [35], a process triggered by multiple factors including energy deficiency [36]. In current study, mitophagy was enhanced in the IF1-KO mice, which is consistent with the phenotypes after IF1 inhibition by shRNA-mediated gene knockdown [25] or gene knockout [37]. However, the role of IF1 in mitophagy remains to established as IF1 knockdown inhibited mitophagy in a cellular model for a deficiency in Parkin (Park2) recruitment [38]. The increase in mitophagy and insulin sensitivity provides a mechanism for the enhanced lipid accumulation in white adipocytes underlying the increased obesity in IF1-KO mice in our study. Mitophagy reduces the mitochondrial number in adipocytes to decrease energy expenditure in support of triglyceride storage [39]. Insulin stimulates triglyceride synthesis and storage in adipocytes. These activities may attenuate the GLP-1 effect in the control of body weight in the IF1-KO mice. The enhanced mitophagy provides a unified mechanism for the enhanced obesity, insulin sensitization and resistance to apoptosis in the IF1-KO mice. Inhibition of autophagy has opposite effects on metabolism as documented in reviews [35; 40]. The IF1 activity is likely mediated by the reduction in ANT2 activity. An enhanced mitophagy is reported in the presence of ANT inhibition by a pharmacological agent [41]. ANT2 reduction may contribute to the control of ATP homeostasis by reducing ADP supply to mitochondria. The mechanism of ANT2 reduction in the IF1-KO mice remains to be investigated.

In summary, the role of IF1 protein is investigated in the pathogenesis of obesity in this study. IF1 protein was reduced in mitochondria by obesity, which represents a compensatory response to the energy surplus in obesity. The conclusion is supported by the metabolic phenotype of IF1-KO mice in both male and female mice, in which obesity was enhanced with a disassociation to insulin resistance. The mitochondrial reprogramming is an underlying mechanism for the enhanced insulin sensitivity and enhanced GLP-1 secretion. The reprogramming was observed in the mitochondrial function and structure in IF1-KO mice. The ATPase activity was enhanced, but ANT2 activity was reduced to limit the ADP supply in mitochondria in the maintenance of ATP homeostasis. The reprogramming was observed with an increase in mitophagy. ANT2 may mediate the IF1 activity in the mitochondrial reprogramming. The molecular mechanism of ANT2 reduction remains to be investigated in the IF1-KO mice.

## Author contribution

Y.W, Y.G, J.Z, X.C, S.S, G.Z, X.X, and X.Z. conducted the experiments. H.W and J.Y designed the study and prepared the manuscript. All authors read and approved the final manuscript.

## Acknowledgment

This study was funded by the National Key Research and Development Program of China (2018YFA0800603 to JY). The authors have no conflict of interest in the publication of this study.

